# The mitochondrial copper chaperone COX11 plays an auxiliary role in the defence against oxidative stress

**DOI:** 10.1101/438101

**Authors:** Ivan Radin, Uta Gey, Luise Kost, Iris Steinebrunner, Gerhard Rödel

**Author notes:** Department of Biology, Washington University in St. Louis, St. Louis, USA. Corresponding authors: (IR) and (GR).

## Abstract

COX11, a protein anchored in the inner mitochondrial membrane, was originally identified as a copper chaperone delivering Cu^+^ to the cytochrome *c* oxidase of the respiratory chain. Here, we present evidence that this protein is also involved in the defence against reactive oxygen species. Quantitative PCR analyses in the model plant *Arabidopsis thaliana* revealed that the level of *AtCOX11* mRNA rises under oxidative stress. The unexpected result that *AtCOX11* knock-down lines contained less ROS than the wild-type can possibly be explained by the impaired oxidative phosphorylation, resulting in less respiration-dependent ROS formation. Similarly, we observed that yeast *Saccharomyces cerevisiae ScCOX11* null mutants produced less ROS than wild-type cells. However, when exposed to oxidative stress, yeast strains overexpressing *ScCOX11* or *AtCOX11* showed lower ROS levels compared with the control indicating a ROS-detoxifying effect of the COX11 proteins. The additive effect on ROS sensitivity upon deletion of *ScCOX11* in addition to the known ROS scavenger gene *SOD1* encoding superoxide dismutase 1 corroborates the oxidative stress-relieving function of ScCOX11. Moreover, yeast strains overexpressing soluble versions of either *AtCOX11* or *ScCOX11* became more resistant against oxidative stress. The importance of three conserved cysteines for the ROS scavenger function became apparent after their deletion that resulted in the loss of ROS resistance. Further studies of strains producing COX11 proteins with individually mutated cysteines indicate that the formation of disulphide bridges might be the underlying mechanism responsible for the antioxidative activity of COX11 proteins. Both AtCOX11 and ScCOX11 apparently partake in oxidative stress defence by directly or indirectly exploiting the redox capacity of their cysteine residues.

## Introduction

For many organisms, aerobic cellular respiration is an essential process, which converts chemical energy stored in sugars and other metabolites into ATP. This complex process is completed by the mitochondrial electron transport chain which shuttles electrons from NAD(P)H and succinate to the terminal acceptor, molecular oxygen [1]. During this process some electrons escape and reduce molecular oxygen, generating superoxide, which can subsequently be converted into other reactive oxygen species (ROS) [2]. While respiratory complexes represent a major source of ROS in mitochondria, several other redox reactions also contribute to ROS production [3]. It is estimated that 1-5% of molecular oxygen is converted to ROS [4].

ROS molecules are highly reactive and can oxidize and thereby damage other molecules such as lipids, proteins, and nucleic acids. Consequently, organisms have evolved complex mechanisms to control ROS levels and reduce their toxicity and detrimental effects (reviewed in [2] and [3]). Some of them are well characterised, for example, the enzyme family of superoxide dismutases (SOD), which convert superoxide ions into oxygen and hydrogen peroxide [5]. The contribution of other proteins to oxidative defence is less well understood and often speculative. One such example is the COX11 (<u>c</u>ytochrome *c* <u>ox</u>idase <u>11</u>) protein family.

Based on data mainly obtained from studies in yeast and bacteria, it is assumed that the main role of COX11 proteins is to deliver Cu^+^ to the Cu_B_ centre of the COX1 subunit of the COX complex (cytochrome *c* oxidase or complex IV of the respiratory chain) [6].

Dimeric COX11 proteins [7] are present in most respiring organisms, from which the homologue of the yeast *Saccharomyces cerevisiae* (ScCOX11) is probably the best-studied family member [8,9,10]. In our previous work, we identified and characterised the *Arabidopsis thaliana* COX11 homologue (AtCOX11) [11]. This homologue is, like the yeast counterpart, localised to mitochondria, presumably to the inner membrane, and involved in COX complex assembly. Interestingly, not only knockdown (KD) but also overexpression (OE) of *AtCOX11* reduced COX complex activity by ~50% and ~20%, respectively [11]. We proposed that both surplus and shortage of COX11 may interfere with the fine-tuned copper delivery balance necessary for COX complex assembly. In line with this, the absence of ScCOX11 leads to a non-functional COX complex and respiratory deficiency in yeast [6,8,12].

However, members of this conserved protein family might be directly involved in mitochondrial oxidative metabolism, as suggested by several publications [13,14,15,16]. Pungartnik *et al.* [13] showed that the yeast *Sccox11* null mutant is highly sensitive to the ROS inducing chemicals N-nitrosodiethylamine and 8-hydroxyquinoline. Subsequently, Khalimonchuk *et al.* [14] and Veniamin *et al.* [15] demonstrated that the *ΔSccox11* strain also showed an increased sensitivity to hydrogen peroxide when compared with the WT strain. For the rice (*Oryza sativa*) COX11 homologue (OsCOX11), direct scavenging of ROS was suggested [16]. The authors reported that OsCOX11 dysfunction leads to a loss of pollen viability, presumably because the timing of a ROS burst necessary for pollen maturation is disturbed. Our previous investigation on AtCOX11 also hinted at its contribution to ROS homeostasis during pollen germination [11]: both the *AtCOX11* KD and OE lines exhibited reduced pollen germination rates, which did not correlate with the observed changes in COX activity, suggesting that AtCOX11 may have an additional function during pollen germination besides COX assembly.

However, the role of COX11 in ROS homeostasis remained elusive. Here, we present our data of a more detailed investigation of COX11’s involvement in oxidative metabolism. Our results indicate that both *Arabidopsis* and yeast COX11 partake in oxidative stress defence, possibly directly by scavenging ROS.

## Material and methods

### Plant material and culture conditions

*Arabidopsis thaliana* (At) Columbia (Col) 0 was used as the WT. The *AtCOX11* knock-down (KD) and overexpressing (OE) lines were previously generated and characterised [11]. KD1/OE lines and KD2 lines were used in T3 and T2 generations, respectively.

Plants were grown either on MS (1× Murashige and Skoog salts, 1% [w/v] sucrose, 0.5 g/L 4-morpholineethanesulfonic acid [MES], 0.8% [w/v] agar) plates or on soil (Einheitserde, type P, Pätzer, Sinntal-Jossa, Germany; mixed with sand 4:1, fertilised by watering with 0.1% [v/v]) Wuxal Basis, Aglukon). For protoplast generation, the MS + 1% sucrose media (for KD lines) was supplemented with 30 μg/mL of kanamycin.

Plants were cultured in a growth chamber with a light intensity of 150 μmol/m^2^s, relative humidity of 35% and day/night temperatures of 24/21°C, respectively. Two types of day/night cycles were used: long day (16-h d) and short day (10-h d).

### Yeast material and culture conditions

*Saccharomyces cerevisiae* (Sc) WT strain BY4741 (Accession (Acc.) number (no.) Y00000) and deletion strains *∆Sccox11* (Acc. no. Y06479) and *∆Scsod1* (Acc. no. Y06913 and Y16913) were obtained from EUROSCARF (Frankfurt, Germany). The *Δcox11Δsod1* strain (MAT *a*; *his3Δ1; leu2Δ0; ura3Δ0; YJR104c::kanMX4; YPL132w::kanMX4*) was generated by crossing the respective single-deletion strains followed by sporulation, tetrad dissection and analysis.

Constructs used for *COX11* overexpression (*pAG415ADH-AtCOX11* and *pAG415ADH-ScCOX11*) were generated previously [11]. To create the soluble versions of COX11, fragments were amplified by PCR (for primer sequences and cloning details see S1 Table) and inserted by Gateway cloning into pDONR or pENTR vectors. All constructs were moved into the high-copy yeast expression-vector pAG425GPD-ccdB-EGFP [17]. Yeast cells were transformed as described in Gietz and Schiestl [18]. Transformed yeast strains were cultured on minimal media (0.5% [w/v] ammonium sulphate, 0.19% [w/v] yeast nitrogen bases, 2% [w/v] glucose, 2.5% [w/v] agar and required amino acids). For oxidative stress tests, YPD (yeast peptone dextrose) media (1% [w/v] yeast extract, 2% [w/v] peptone, 2% [w/v] glucose, 2% [w/v] agar) supplemented with the corresponding oxidative stressors was used. Media were cooled to 55°C; freshly prepared chemical stocks were added just before pouring, and plates were used within 24 h. For liquid cultures, yeast strains were cultured at 30°C with shaking at 180 rpm.

For growth analysis, yeast strains were cultured in liquid minimal media for 24 h, then diluted with minimal media to OD_600_ = 0.05 and cultured for another 16 h. Serial dilutions were spotted on solid media plates (YPD with or without oxidative stressors). Growth was documented after incubation for 48-60 h at 30°C.

### Bioinformatic analysis

*Arabidopsis* and yeast gene and protein sequences were obtained from The *Arabidopsis* Information Resource [19] and the GeneBank [20], respectively. For protein sequence alignment, the EMBOSS Needle software (The European Bioinformatics Institute) [21] was used. For the prediction of targeting signal cleavage sites, TargetP [22,23] was used, and the transmembrane domains were predicted with TMHMM2.0 [24]. Disulphide bridge formation in proteins was predicted with DiANNA 1.1 [25,26,27]. The Genevestigator was used to examine public microarray databases [28].

### Stress treatments and qPCR

For the oxidative stress treatments, the *Arabidopsis* WT seedlings were cultured on solid MS plates + 1% (w/v) sucrose for 12 days. Stress was applied for 2 h or 6 h by placing seedlings on the surface of liquid MS + 1% (w/v) sucrose media supplemented with the appropriate stressor. Antimycin A (Sigma Aldrich) stock was dissolved in absolute ethanol and subsequently diluted with MS media. As a control, seedlings were placed on the surface of liquid MS + 1% (w/v) sucrose media without the stressors. Immediately after the stress treatment, the seedlings were frozen in liquid nitrogen, and RNA was isolated. RNA isolation and quantitative real-time RT-PCR (qPCR) were performed as previously described [11]. The RNA quality was analysed with the BioAnalyzer 2100 (Agilent, USA), and only RNAs with RNA integrity numbers (RIN) in the range of 7.5 to 8.5 were reverse transcribed. The efficiency and optimal concentrations of all primer pairs were experimentally determined and are listed in the S2 Table. The data were statistically analysed with the Bio-Rad CFX Manager 3.1 software.

### Lipid peroxidation measurement

The levels of lipid peroxidation were determined with the Bioxytech LPO-586 kit (OxisResearch, USA). Rosette leaves from plants (10 weeks old) grown under short-day conditions were harvested at the beginning of the light period and immediately ground with a pestle in a mortar with 500 μL of grinding buffer (20 mM Tris-Cl pH 7.4, 5 mM butylated hydroxytoluene) per 100 mg of tissue. The leaf suspension was cleared by two centrifugation steps (each 3,000*g* for 10 min at 4°C). Of the final supernatant, 7 μL and 100 μL were used for quantitation of protein concentration (Bio-Rad DC assay, USA) and lipid peroxidation measurements, respectively. The “reagent 2” (methanesulfonic acid) was employed to determine the amounts of malondialdehyde (MDA) and 4-hydroxyalkenals (HAE). All samples were run in triplicates and read out with a TECAN M200 plate reader (Tecan, Switzerland). The lipid peroxidation levels were normalised to the protein concentrations in the supernatants.

### ROS level measurement in protoplasts

Protoplasts were isolated as previously described [29] with slight modifications. Of the protoplasting buffer (20 mM KCl, 20 mM 4-morpholineethanesulfonic acid [MES], 0.4 M mannitol, 1.25% [w/v] cellulase R-10, 0.3% [w/v] macerozyme R-10, 10 mM CaCl_2_, 0.1% [w/v] BSA, pH 5.7) 1.5 mL were added to approximately 100 mg of finely cut 12-d-old seedlings cultured under long-day conditions. After 4 h of agitation at room temperature, the suspensions were successively filtered through 100- and 50-μm meshes. Protoplasts were pelleted (280*g* for 10 min at 4°C) and washed first with W5 buffer (154 mM NaCl, 125 mM CaCl_2_, 5 mM KCl, 5 mM MES, pH 5.7), and then with MMG buffer (0.4 M mannitol, 15 mM MgCl_2_, 4 mM MES, pH 5.7). They were finally resuspended and stored in MMG buffer at 4°C until use. All buffers were prepared fresh.

For determination of ROS levels, protoplasts were incubated with 5 μM DCFDA (2’,7’ dichlorofluorescin diacetate) for 10 min [30] and then imaged with the LSM780 microscope from Zeiss (C-Apochromat 40×/1.20 W Korr M27 objective, excitation with 488-nm laser, detection in 510-542 nm range for DCF (2’,7’ dichlorofluorescin) and 647-751 nm for chlorophyll autofluorescence). The total fluorescence was determined with the Fiji image analysis software [31] as raw integrated density in the green channel of individual protoplasts. Protoplasts with chloroplasts, which are derived from photosynthetic tissues, were excluded to avoid measurement of ROS produced by photosystems.

### ROS level measurement in yeast cells

Liquid YPD media was inoculated with the respective strains and cultured for 24 h. Then the cultures were diluted to OD_600_ of 0.01, grown overnight (14-16 h) and used to start the final YPD cultures (starting OD_600_ = 0.1), which were incubated at 30°C until an OD_600_ of 0.5-0.6 was reached. This successive refreshing was necessary to ensure the same physiological state of all strains. Cultures were aliquoted (1 mL each) into 2-mL tubes and either treated with water (= mock) or with 2 mM paraquat (PQ; methyl viologen from Sigma Aldrich) for 30 min at 30°C with agitation. Subsequently, cells were pelleted (3500*g* for 3 min at RT) and washed twice with PBS. Finally, cells were resuspended in 1 mL of PBS and split into two aliquots. One was used as the negative control, while the other was stained with DCFDA (final concentration 20 μM) for 45 min at 30°C with agitation. After staining, cells were washed twice with PBS. Total DCF fluorescence was measured in the CyFlow SL (Partec, Germany) with 488-nm excitation and detection in FL1 channel (527 nm/BP 30 nm). FL1 channel gain was set to a level on which fluorescence could not be observed in negative unstained controls.

## Results

### Oxidative stress induces *AtCOX11* expression

As a first step to investigate a role of *AtCOX11* in ROS homeostasis, as previously proposed [11], we analysed its promoter region for the presence of *cis*-active ROS-responsive elements which are prevalent in known ROS-induced genes [32,33]. In *AtCOX11*, the non-coding region upstream of the start codon harbours as many as 16 putative oxidative-stress-responsive elements (Fig 1A and S1 Fig). In contrast, the promoter region of *AtHCC1*, another mitochondrial chaperone delivering copper to the COX complex [34], contains only five ROS-responsive consensus sequences. AtHCC1 stands for <u>h</u>omologue of <u>c</u>opper <u>c</u>haperone SCO<u>1</u> (<u>s</u>ynthesis of <u>c</u>ytochrome <u>*c*</u> oxidase 1).

**Fig 1.**
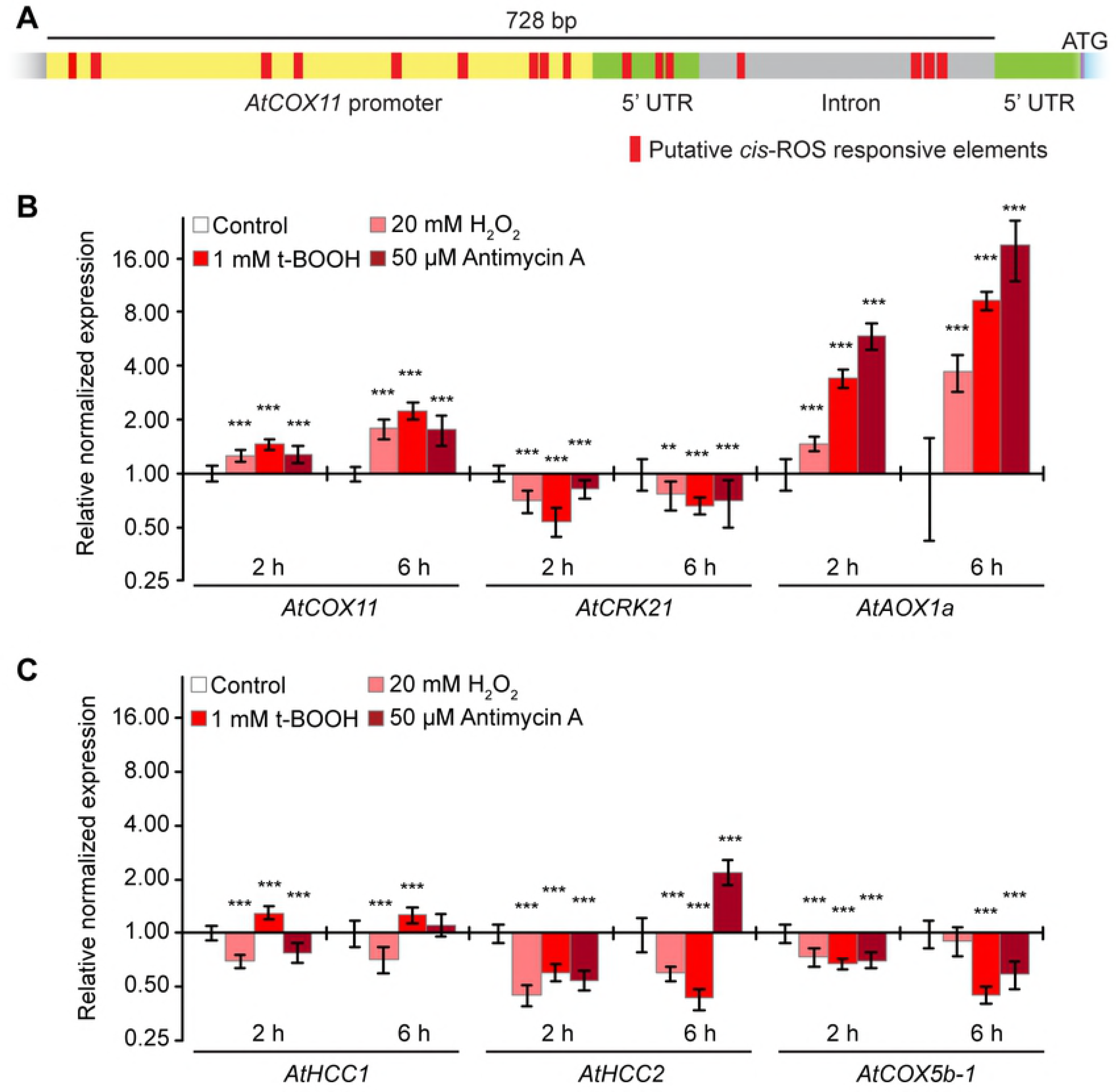
*AtCOX11* is upregulated by oxidative stress. (A) Scaled diagram of putative ROS-responsive elements within the *AtCOX11* promoter and 5’UTR. (B) and (C) Gene regulation in response to oxidative stress. Stress was applied for 2 h and 6 h as described in the methods section. Mean values of mRNA levels in treated samples were normalized to the control sample and plotted on a logarithmic scale (base 2). Values and statistical significance compared with the control sample (**P < 0.01; ***P < 0.001) were calculated with the CFX manager software. Error bars represent ± standard deviation (SD). Individual values and SD are listed in the S3 Table.

The overrepresentation of putative ROS-responsive elements prompted us to analyse the expression levels of *AtCOX11* transcripts under oxidative stress (Fig 1B). We treated WT seedlings for 2 h or 6 h with the oxidative reagents hydrogen peroxide (H_2_O_2_), tert-butyl hydroperoxide (t-BOOH) and antimycin A followed by qPCR analyses. Hydrogen peroxide is a ROS molecule which easily transverses membranes and induces oxidative stress throughout the whole cell, while the organic peroxide t-BOOH is transported to mitochondria as well as other cellular compartments [35]. Antimycin A induces ROS production at the mitochondrial electron transport chain by inhibiting the respiratory complex III [36]

*AtCRK21* (<u>c</u>ysteine-<u>r</u>ich <u>r</u>eceptor-like protein <u>k</u>inase <u>21</u>) and *AtAOX1a* (<u>a</u>lternative <u>ox</u>idase <u>1a</u>) are known to be down- and upregulated [37] by ROS, respectively, and were therefore chosen as controls. As expected, oxidative stress reduced *AtCRK21* levels, while *AtAOX1a* mRNA abundance was increased about 19-fold (Fig 1B).

*AtCOX11* was slightly upregulated (~1.3 fold) in response to all three 2-h oxidative stress conditions. The upregulation further increased to ~2 fold after 6 h. These data suggest that at least some of the regulatory elements present in the *AtCOX11* promoter region are functional.

In order to check whether the *AtCOX11* ROS-response profile is unique and specific, the expression of other COX assembly and subunit genes was analysed under oxidative stress. The transcript levels of the copper chaperone *AtHCC1* were only marginally affected at both time points (Fig 1C). On the other hand, its homologue *AtHCC2* (<u>h</u>omologue of <u>c</u>opper <u>c</u>haperone SCO<u>2</u>), which lacks a copper-binding motif [38], was affected by ROS. It showed a decrease of the transcript level by about half (Fig 1C), even though its promoter region carries seven putative ROS-responsive elements (S1 Fig). *AtHCC*2 levels, 2.2 times higher after a 6-h antimycin A treatment, were a notable exception from the otherwise observed downregulation. Although the *AtHCC2* expression pattern was different from *AtCOX11*, the fact that *AtHCC2* responded to ROS fits a previously proposed role of *AtHCC2* in redox homeostasis [38,39].

The transcript levels of another COX-related gene, the COX subunit *AtCOX5b-1*, were reduced by ~30% after 2 h of oxidative stress and by ~50% after 6 h, except for the H_2_O_2_ treatment, which had no effect at this time point (Fig 1C). Clearly, not all mitochondrial genes respond to ROS, and if they do not in the same way. Our qPCR data for all genes analysed are backed up by public microarray data (Genevestigator database) of a 3-h treatment with 50 μM antimycin A (applied by spraying) [40] (S4 Table).

Taken together, *AtCOX11* shows a unique ROS response characterized by an accumulation of transcripts for all three oxidative stressors applied. In addition, the response increased over time supporting a role of *AtCOX11* in ROS homeostasis as suggested by the enrichment of ROS-responsive elements in its promoter region.

### Knockdown of *AtCOX11* reduces cellular ROS

To explore a role in ROS homeostasis further, ROS levels were measured in the *Arabidopsis COX11* KD and OE plant lines that were generated previously [11]. The *AtCOX11* mRNA levels in KD plants were approximately 30% of the WT levels, while in the two overexpression lines OE1 and OE2, the transcript amounts were approximately 6- and 4-fold higher, respectively [11]. ROS levels were measured by two independent methods: indirectly by determining the lipid peroxidation levels (Fig 2A), and directly by staining protoplasts with the ROS-specific dye DCFDA (Fig 2B). For lipid peroxidation measurements, plants were grown for 14 h in the dark prior to the experiments to minimise ROS contributions from photosystems. Then, the leaves were harvested to measure MDA and HAE concentrations, typical products generated by decomposing lipid peroxides.

**Fig 2.**
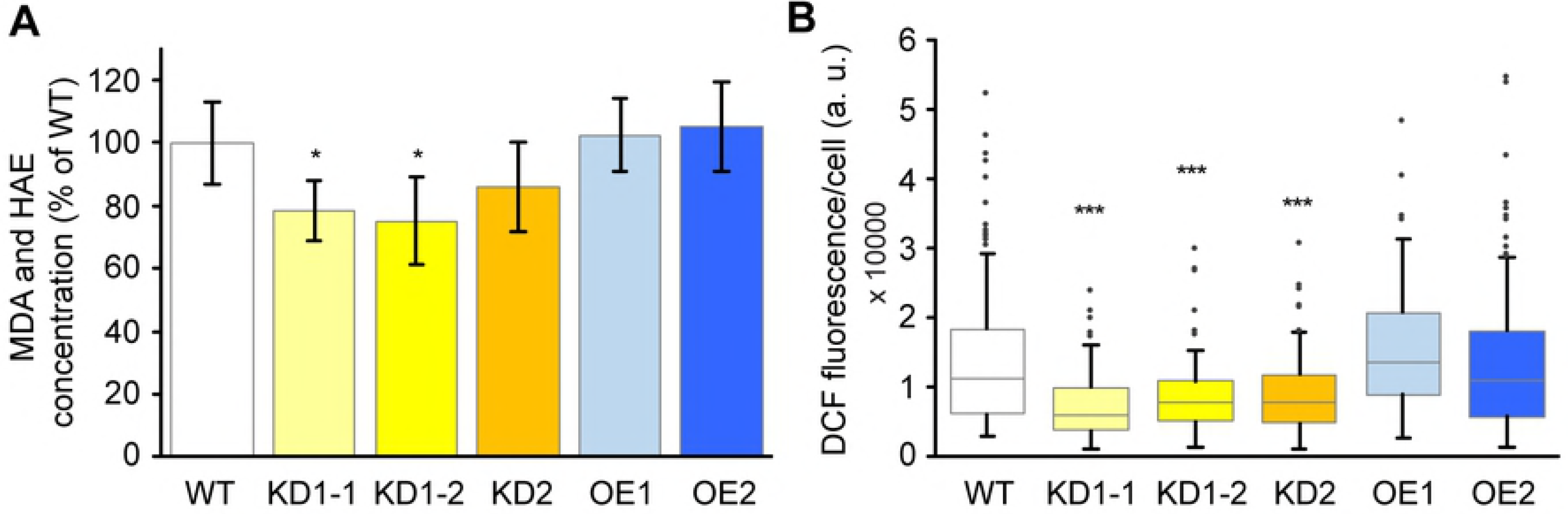
Disturbance of *AtCOX11* expression alters cellular ROS levels. (A) Lipid peroxidation was determined in *AtCOX11* knock-down (KD) and overexpression (OE) mutants by measuring the concentration of malondialdehyde (MDA) and hydroxyalkenals (HAE) in their leaves and normalised to the WT (= 100%). Each bar represents the mean ± SD of five independent experiments. (B) Box plots of DCF fluorescence in arbitrary units (a. u.) as an indicator of ROS levels in protoplasts from WT and *AtCOX11* KD and OE mutants are shown. The distributions of fluorescence intensities of individual protoplasts from various genotypes are depicted. For each box the horizontal lines designate the median, and first and third quartile. The vertical lines and dots extending from each box mark the lowest and the highest fluorescence values. Asterisks indicate statistically significant difference (unpaired Student’s t-test; *P < 0.05, ***P < 0.001) between mutants and WT. The absolute and normalised values for (A) and descriptive statistics for (B) are given in S3 Table.

MDA and HAE levels were lower in all KD lines compared with the WT, albeit only statistically significant for KD1-1 and KD1-2 plants (Fig 2A). The levels in the OE lines were indistinguishable from the WT.

These data were confirmed by a second assay, in which protoplasts were incubated with the DCFDA dye, which upon entering the cell and oxidation by ROS exhibits a bright green fluorescence. All KD lines showed a statistically significant reduction in cellular ROS levels compared with the WT and again, the OE lines were indistinguishable from the WT (Fig 2B). Of note is that these assays detect ROS from the entire cell and might not be sensitive enough to detect subtle changes in the intermembrane space of mitochondria.

These results seemingly contradict a function of *Arabidopsis* COX11 in ROS defence. However, the observed phenotypes in the KD lines could be contributed to the loss of COX complex activity (see discussion for details). In summary, two different ROS detection methods revealed a reduction in ROS levels when *AtCOX11* expression was reduced, but no change in ROS amounts when *AtCOX11* was overexpressed.

### COX11 proteins play a role in oxidative stress tolerance in yeast

Next, we investigated the role of COX11 proteins in ROS homeostasis in another model organism, the budding yeast (*S. cerevisiae*). For this, *ScCOX11* was knocked out or overexpressed (alternatively *AtCOX11)* and the effects on cellular ROS levels were studied under normal and oxidative stress conditions (Fig 3A and 3B). Yeast cells were stained with the ROS-specific dye DCFDA, and the green fluorescence of each cell was measured by flow cytometry. For each data set, the mode, defined as a number that occurs most often in the data set, was determined. Mode corresponds to the X-axis position of the peak of the cell fluorescence intensity histogram (S2 Fig). Modes from three independent experiments were averaged and depicted as bar graphs (Fig 3A and 3B). Cell fluorescence intensity histograms representing the data from individual experiments are depicted in S2 Fig.

**Fig 3.**
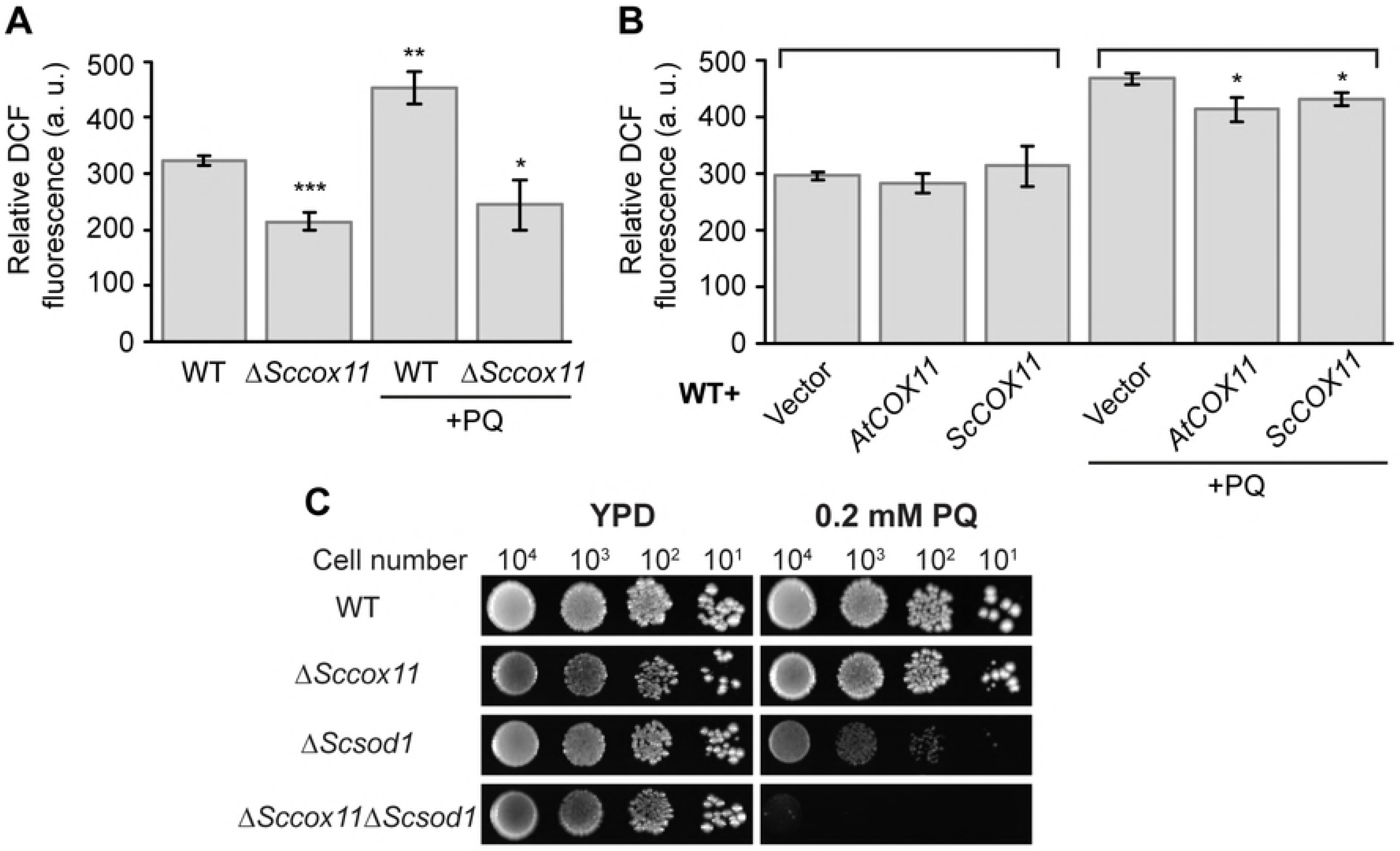
COX11 proteins influence yeast oxidative stress tolerance. ROS levels determined by DCFDA staining of WT, *ΔSccox11* (A) and *AtCOX11* or *ScCOX11* overexpressing (B) yeast strains; after either mock treatment or treatment with 2 mM paraquat (PQ). Total fluorescence of individual cells was measured by flow cytometry. Descriptive statistics for the cytometry datasets are given in S3 Table. Graphs in (A) and (B)depict averages of modes (see text for details). Each bar represents the mean of modes ± SD from three independent experiments. Asterisks indicate statistically significant difference (unpaired Student’s t-test; *P < 0.05, **P < 0.01, ***P < 0.001) between mode averages of mutant strains compared with the untreated WT (A) or corresponding empty vector control of the treated or untreated dataset (B) under the same treatment. (C) Growth on normal and oxidative stress media, of *ScCOX11* and *ScSOD1* double-deletion yeast strain was compared with single-deletion mutants as well as the WT strain.

Like in plants (Fig 2), the *ScCOX11* knock-out (KO; *ΔSccox11*) strain showed a significant reduction in the cellular ROS levels compared with the WT strain (Fig 3A). The overexpression of either the yeast or plant COX11 protein did not affect ROS levels compared with the control strain transformed with the empty vector (Fig 3B). In addition, we treated all strains with 2 mM PQ to test whether the KO or OE of *COX11* changes tolerance to oxidative stress. PQ is a ROS inducer and a redox cycler that targets primarily electron transport chains (ETC), [41,42].

PQ significantly increased cellular ROS levels in the WT compared with the untreated control (Fig 3A). The same treatment did not affect ROS levels in the respiratory deficient *ScCOX11* KO strain (Fig 3A). The *AtCOX11* or *ScCOX11* overexpressing yeast strains showed increased ROS levels in response to PQ (Fig 3B). However, the ROS levels’ increase was slightly, but significantly smaller compared with the increase in the empty-vector control (Fig 3B). This indicates that the overexpression of COX11 genes can partly alleviate the oxidative stress. The reduction in ROS levels in the intermembrane space (IMS) might even be higher, because the DCFDA-staining results from total cellular ROS, thereby possibly masking the small contributions of the mitochondrial IMS compartment.

An intriguing possibility might be that the role of ScCOX11 in ROS defence is redundant with main ROS defence mechanisms such as the action of ScSOD1, which is localised both in the cytoplasm and IMS [43]. To test this hypothesis, a *ScCOX11* and *ScSOD1* double-deletion mutant (*ΔSccox11ΔScsod1*) was generated by crossing the respective single-deletion strains. The growth of the WT, double-deletion and the corresponding single-deletion strains was analysed under standard conditions (YPD) or oxidative stress (YPD + 0.2 mM PQ) (Fig 3C). On YPD media, all strains showed comparable growth. The added PQ did not affect the growth of the WT and *ΔSccox11* mutant. As expected, the *ScSOD1* (*ΔScsod1*) single-deletion strain showed a strong reduction in growth. Strikingly, the growth of the double-deletion mutant *ΔSccox11ΔScsod1* was even more severely reduced, indicating partially overlapping functions of COX11 and SOD1.

Taken together, these results suggest that COX11 proteins are not major players of the ROS defence mechanism, but might contribute to ROS detoxification, possibly by scavenging.

### Soluble COX11 proteins improve yeast growth under oxidative stress

The big challenge in studying oxidative stress is the sensitivity of ROS detection assays. COX11 proteins function in the mitochondrial IMS, which is a rather small compartment. A change in ROS levels may be, at least partially, hidden due to ROS contributions from other cellular compartments. To circumvent this problem, we took another approach to test the ROS protective ability of COX11 proteins. Constructs were generated expressing soluble (sol) versions of the *Arabidopsis* and yeast COX11 proteins (Fig 4A) lacking the mitochondrial targeting signals and almost the entire transmembrane domains (TM) except for seven highly conserved amino acids (Fig 4A and S3 Fig).

**Fig 4.**
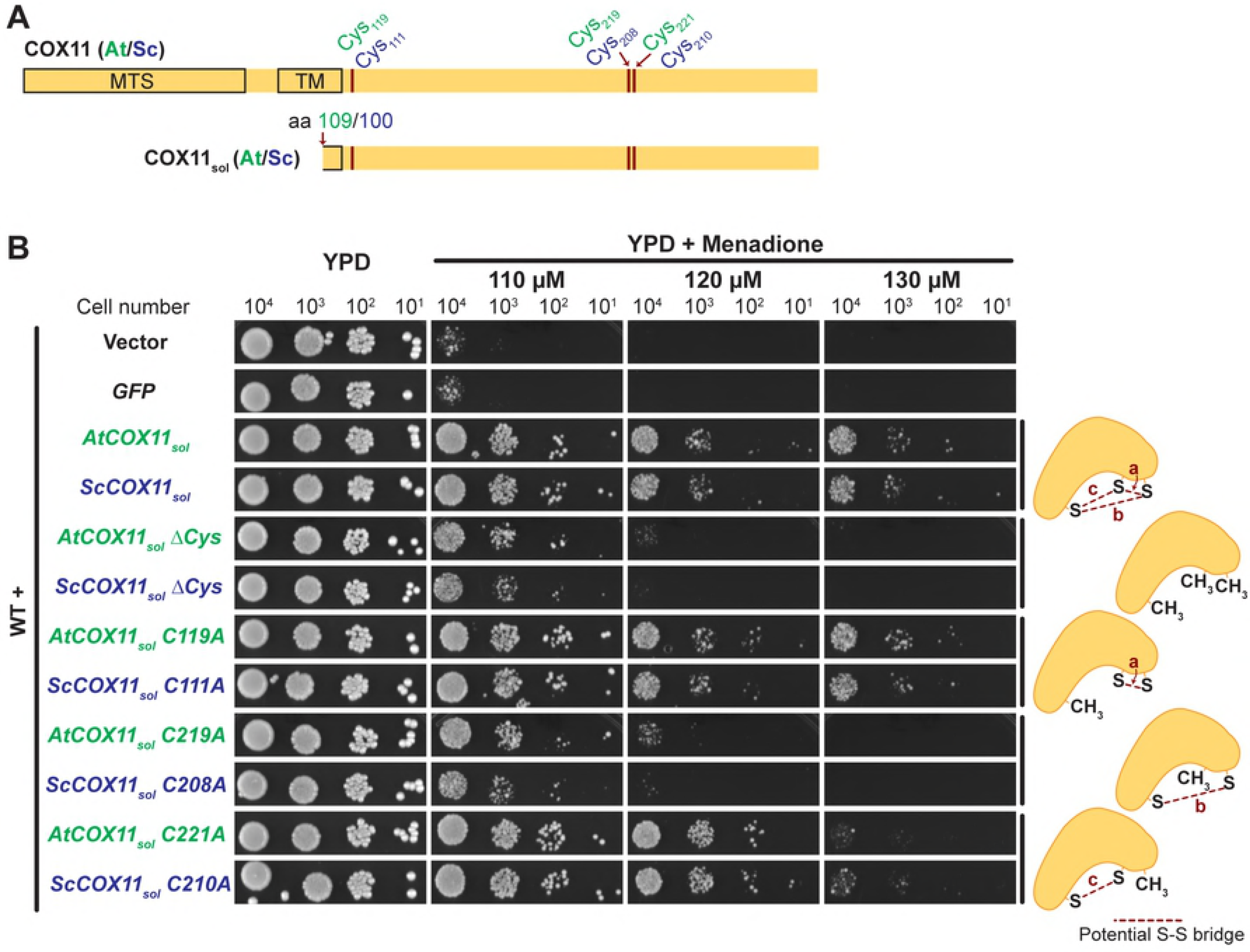
Soluble COX11 proteins improve yeast growth under oxidative stress. (A) Scaled diagram of full-length COX11 proteins from *Arabidopsis thaliana* and *Saccharomyces cerevisiae* (top scheme) and truncated versions (AtCOX11_sol_ = AtCOX11 without amino acids (aa) 2-108; ScSCOX11_sol_ = ScCOX11 without aa 2-100) (bottom scheme). The protein alignment and domain details are given in S3 Fig. The positions of cysteines (Cys) are indicated. aa (amino acids), MTS (mitochondrial targeting signal), TM (transmembrane domain), sol (soluble). (B) The growth of WT yeast expressing different soluble versions of *Arabidopsis* or yeast COX11 proteins under oxidative stress. In the mutated soluble COX11 versions, either all three cysteines (ΔCys) or only one was replaced with an alanine. Serial dilutions of yeast strains were spotted on YPD media without or with increasing concentration of the cellular oxidative stressor menadione. Diagrams on the right visualize possible disulphide-bridge formation in each protein version (see text for details). Depicted growth assays are exemplary from at least three independent experiments.

Control constructs consisted of the empty expression vector and the vector expressing green fluorescence protein (GFP). GFP has a similar molecular weight as COX11, so its expression should exert the same energetic cost on the cells as the expression of *COX11*, making it a suitable control. WT yeast cells were transformed with these constructs and their growth monitored on YPD plates (Fig 4B). Menadione was chosen as the oxidative stressor because it is a known general redox cycler and ROS inducer in the cytoplasm and other compartments [44].

All yeast strains grew equally well in the absence of oxidative stress. When menadione was added to the medium, however, the empty-vector, as well as the GFP-expressing controls, were almost unable to maintain growth, even at the lowest menadione concentration (Fig 4B). The halted growth of the GFP control shows that the overexpression of a random protein does not confer oxidative stress tolerance.

The yeast strains expressing either *AtCOX11*_*sol*_ or *ScCOX11*_*sol*_, however, continued to grow at all three menadione concentrations tested (Fig 4B). At the lowest concentration of 110 μM, growth remained almost unaffected. These results indicate that the increased menadione tolerance in yeast expressing soluble COX11 is likely linked to some intrinsic feature(s) of the COX11 proteins.

What is the feature that allows COX11 proteins to heighten resistance to oxidative stress? One possibility would be the three highly conserved cysteines present in COX11 proteins, of which two belong to the copper-binding motif (Fig 4A and S3 Fig) [7]. There are additional cysteines present in the N-termini of the COX11 proteins (S3 Fig), but they are part of the predicted mitochondrial targeting signal and therefore absent in the mature proteins.

To test the importance of the conserved cysteines, we generated the mutant strain Δcys in which the three cysteines were converted into alanines. This strain was still able to moderately grow in the presence of 110 μM menadione, but not of 120 and 130 μM. This result demonstrated that the conserved cysteines apparently do play a role in the ability of COX11 proteins to diminish the oxidative stress burden, possibly by directly detoxifying ROS molecules through oxidation and formation of intracellular disulphide bridges.

To find out which of the three possible bridges (labelled “a”, “b” and “c” in the schematic illustrations in Fig 4B) might be involved, we generated six more constructs, three *Arabidopsis* and three yeast *COX11* versions, in which in each case one of the three cysteines was mutated to an alanine thus restricting the number of putative disulphide bridges that can be formed (illustrated in the schemes on the right of Fig 4B). The yeast strains transformed with the *COX11* versions that could either form bridge “a” or bridge “c” retained their capacity to improve the resistance of the cells to oxidative stress, similar to the strains expressing the soluble versions with all three cysteines. In contrast to that, the strains expressing the versions which could only form bridge “b” showed the same reduced stress resistance as the Δcys versions. These results suggest that the formation of either bridge “a” (Cys_208_ and Cys_210_ in yeast; Cys_219_ and Cys_221_ in *Arabidopsis*) or bridge “c” (Cys_111_ and Cys_208_ in yeast; Cys_119_ and Cys_219_ in *Arabidopsis*) or both might be the feature that allows COX11 proteins to detoxify ROS.

In summary, our data show that soluble forms of COX11 proteins increase the oxidative stress tolerance in yeast involving the conserved cysteines, possibly through the formation of disulphide bridges.

## Discussion

The role of COX11 proteins as copper chaperones in COX complex assembly has been well documented [8,9,11,12,45]. In this work, we present evidence that COX11 proteins have an auxiliary role in the defence against oxidative stress.

The initial hint for such a role came from our observation that the expression of the *Arabidopsis COX11* gene was upregulated in response to oxidative stress (Fig 1B). This appeared to be a specific response of the *AtCOX11* gene and not part of a general upregulation of mitochondrial genes because *AtHCC1* levels, for example, remained unchanged and *AtHCC2* and *AtCOX5b-1* genes were downregulated (Fig 1C). Interestingly, *AtHCC2*, which has also been implicated in ROS defence after UV-B light exposure [38], responded to the chemical oxidative stressors mostly with downregulation. When antimycin A was applied, however, the *AtHCC2* transcript levels were initially reduced but increased after 6 h (Fig 1C). These findings confirm previous reports on the sensitivity of the oxidative defence machinery to the type of stressor and the time point of analysis [37,46]. Taken together, this data supports that AtCOX11 likely has an auxiliary role in the oxidative defence in addition to its main role in copper transport.

One would expect that knockdown and overexpression of an oxidative stress defence protein to result in higher and lower ROS levels, respectively. Nevertheless, at first glance, our experiments did not fulfil these predictions and even yielded opposite results with knock-down plant mutants having reduced ROS levels (Fig 2). This reduction could be the result of the lower COX complex activity found in these plants (~50% of the WT [11]), as previously reported in mice mutants, where COX deficiency led to decreased oxidative stress [47]. The absence of a functional COX has repeatedly been reported to result in the downregulation of other respiratory complexes [48,49], eventually reducing the ROS load that is typically associated with the functional respiratory chain [4]. Moreover, the increased expression of alternative oxidases in COX-deficient mutants, as observed in *AtCOX11* KD plants [11], is probably a mechanism to compensate for the COX loss and was shown to lower mitochondrial reactive oxygen production in plant cells [50]. Therefore, both the COX deficiency and the expression of alternative oxidases may mask the reduced ROS-scavenging contribution of COX11 in the *AtCOX11* KD lines.

Unexpectedly, no difference in total cellular ROS amounts was found between *COX11* OE plants and the WT. However, as the mitochondrial IMS accounts for only a minor portion of the ROS-producing cellular compartments, a possible ROS-scavenging effect by a mild *AtCOX11* overexpression may have escaped detection.

Analogous experiments with yeast *ScCOX11* knock-out and overexpressing strains cultured under standard conditions yielded similar results as observed in plants (Figs 2 and 3): lower ROS levels in *ΔSccox11*, and levels indistinguishable from the WT in the overexpressing strains. However, when the *COX11* OE strains were treated with PQ, their ROS levels were lower compared with the treated empty-vector control (Fig 3B and S2 Fig). Therefore, it appears that COX11 proteins confer some level of protection under oxidative stress conditions. Alternatively, the difference between ROS levels was large enough to be detected in this experimental setup.

Further evidence that COX11 proteins are involved in mitochondrial oxidative defence came from the *ScCOX11* and *ScSOD1* double-deletion strain (Fig 3C). The fact that this strain showed a much higher sensitivity to PQ than either single-deletion or WT strains indicates that ScCOX11 and ScSOD1, both of which function in the IMS, have overlapping and additive functions. COX11 proteins might help the main mitochondrial ROS defence players, like SOD1, under heightened oxidative stress or even normal conditions. The COX11 proteins, as COX complex chaperones, are in the vicinity of ROS-generating respiratory complexes and could therefore potentially quickly detoxify ROS and prevent damage.

These results suggest that COX11 proteins might directly or indirectly affect ROS levels in the IMS. However, as already mentioned, evaluation of ROS levels in the mitochondrial IMS is technically challenging. Therefore, we generated genetic constructs for the expression of soluble versions of AtCOX11 and ScCOX11 in the cytoplasm of yeast cells (Fig 4A). Both soluble versions permitted growth in the presence of ROS-inducing menadione (Fig 4B), showing that COX11 proteins are indeed able to reduce oxidative stress. Since the antioxidative function was exerted even in the non-native cellular environment, one may speculate that COX11 proteins are able to function in ROS defence, independently of other proteins.

The mutation of all three cysteines in the COX11 proteins mostly abolished their ability to convey growth under oxidative stress, emphasizing the role of these amino acid residues in ROS detoxification. However, when compared with the GFP-expressing control strain, the triple cys mutants were still more resistant to menadione, hinting at an additional ROS-protective mechanism aside from cysteine oxidation. For example, the oxidation of other COX11 amino acids side chains (e.g. methionine; S3 Fig) by ROS molecules. Based on the data from the various mutants, the most fitting mechanism is that COX11 is scavenging ROS directly by the formation of disulphide bridges (S-S) between the conserved cysteines (Fig 4B, see diagrams on the right) as previously reported for other ROS protectants, e.g. the human PRX3 (peroxiredoxin-3) [3]. To address this hypothesis, we analysed the contribution of the cysteines and putative S-S bridges between them (named a, b and c in Fig 4B) to the observed COX11 antioxidant activity. We generated variant forms of COX11 with individual cysteines mutated to alanines, only allowing the formation of a single putative S-S bridge (Fig 4B, right). The cysteine combinations 119/219 or 219/221 in *Arabidopsis* and 111/208 or 208/210 in yeast, maintained growth under oxidative stress. Interestingly, the various menadione concentrations used, highlight the sensitivity of oxidative stress tests. At a concentration of 110 μM the loss of one of the three cysteines had no effect, but a mere increase of 10% to 120 μM made the difference in oxidative stress resistance readily apparent. Specifically, the loss of cysteine 219/208 (*Arabidopsis*/yeast) diminished the antioxidant activity, corroborating the hypothesis that the ability to form the S-S bridges a and/or c is crucial.

In their study of yeast COX11, Bode *et al*. [51] provided experimental evidence for the formation of the disulphide bridge between the two cysteines 208 and 210 within the Cu-binding motif (= bridge a). On the other hand, the disulphide bond prediction software DiANNA 1.1 predicted only the formation of bridge c, albeit with a low probability score (*Arabidopsis*/yeast COX11 bridge a: 0.01/0.01, bridge b: 0.01/0.01 and bridge c: 0.12/0.16; maximum score: 1). Furthermore, both the previously published crystal structure [52] of a bacterial COX11 (*Sinorhizobium meliloti*) and the model [53] of human COX11 revealed that all three conserved cysteines are on the protein surface and thus easily accessible to oxidation by ROS molecules and S-S bridge formation. Of note is that COX11 proteins - in addition to the formation of intramolecular disulphide bridges within a single COX11 subunit - could potentially also form intermolecular bridges between two COX11 subunits or between COX11 and another protein or e.g. glutathione (GSH).

As an alternative explanation for their antioxidant activity, the COX11 copper chaperones may use the bound copper to detoxify ROS. However, this scenario seems unlikely, because our experiments demonstrate that the loss of one of the cysteines in the copper-binding motif (cys 221 and 210 in *Arabidopsis* and yeast COX11, respectively) did not eliminate the COX11 antioxidant activity (Fig 4B).

Taken together, the findings that the mutation of the respective cysteines had the same positive or negative antioxidant effects in two evolutionary distant organisms like *Arabidopsis* and yeast, pinpoint that these cysteines and their functions were obviously important to be conserved during evolution.

Therefore, it seems plausible that the formation of either disulphide bridge a or c, or both, is the mechanism by which COX11 proteins detoxify ROS. These potentially ROS-induced S-S bridges could subsequently be reduced in the IMS by thioredoxins or proteins with a putative thioredoxin domain such as AtHCC2 [38], or by other redox systems, e.g. the ERV1/MIA40 IMS protein import system [3]. While many open questions remain regarding the role of COX11 proteins in ROS metabolism, the data presented here show that the *Arabidopsis* and *S. cerevisiae* COX11 proteins are able to relieve oxidative stress. COX11 proteins might play a role in alleviating ROS stress generated by the respiratory complexes or are needed under elevated oxidative stress as the second line of defence.

## Acknowledgments

We thank the Light Microscopy Facility of the BIOTEC/CRTD at Technische Universität Dresden for their help with our confocal microscopy experiments.

## Supporting information

**S1 Fig. Cis-acting putative ROS-response elements in the putative promoter regions of *AtCOX11*, *AtHCC1* and *AtHCC2*.**

**S2 Fig. Total cell fluorescence intensity distributions of *ScCOX11* knock-out (A, B, C) and overexpressing yeast (D, E, F) cells stained with DCFDA.**

**S3 Fig. Alignment of *A. thaliana* (A. t.) and *S. cerevisiae* (S. c.) COX11 protein sequences.**

**S1 Table. Cloning primers**

**S2 Table. Primers used for qPCR.**

**S3 Table. Absolute and normalized values from bar graphs and descriptive statistics.**

**S4 Table. Gene regulation in response to 50 μM antimycin A (microarray data from Ng *et al.* [2013], Genevestigator).**

